# Proteomics and phosphoproteomics profiling of the co-formulation of type I and II interferons, HeberFERON, in the glioblastoma-derived cell line U-87 MG

**DOI:** 10.1101/2022.10.03.510562

**Authors:** Dania Vázquez-Blomquist, Anette Hardy-Sosa, Saiyet C. Baez, Vladimir Besada, Sucel Palomares, Osmany Guirola, Yassel Ramos, Jacek R. Wiśniewski, Luis Javier González, Iraldo Bello-Rivero

## Abstract

HeberFERON is a co-formulation of Interferon (IFN)-α2b and IFN-γ in synergic proportions, with a demonstrated effect on skin cancer and other solid tumors. It has antiproliferative effects over glioblastoma multiform (GBM) clones and cell lines in culture, including U-87 MG. Omics studies in U-87 MG showed distinctive expression patterns compared to individual IFNs. Kinase signaling pathways dysregulation can also contribute to HeberFERON effects. Here, we report the first label-free quantitative proteomic and phosphoproteomic analyses to evaluate changes induced by HeberFERON after 72h incubation of U-87 MG cell line. LC-MS/MS analysis identified 7627 proteins with a fold change >2 (p<0.05); 122 and 211 were down- and up-regulated by HeberFERON, respectively. We identified 23549 peptides (5692 proteins) and 8900 phosphopeptides, 412 of these phosphopeptides (359 proteins) were differentially modified with fold change >2 (p<0.05). Proteomic enrichment analysis showed IFN signaling and its control, together to direct and indirect antiviral mechanisms were the main modulated processes. Enrichment analysis of phosphoproteome pointed to the cell cycle, cytoskeleton organization, translation and RNA splicing, autophagy, and DNA repair as biological processes represented. There is a high interconnection of phosphoproteins in a molecular network, where mTOR occupies a centric hub. HeberFERON regulates many phosphosites newly reported or with no clear association to kinases. Of interest is phosphosites increasing phosphorylation were mainly modified by CDK and ERK kinases, thus new cascades regulations can be determining the antiproliferation outcome. Our results contribute to a better mechanistic understanding of HeberFERON in the context of GBM.

**Significance of the Study:** HeberFERON is a co-formulation of IFN-α2b and -γ in synergic proportion, registered for skin basal cell carcinoma treatment, also demonstrating clinical effect over solid tumors, including GBM. GBM is a very lethal tumor, protected by the blood-brain barrier (BBB), highly mutated in proliferative signaling pathways with little treatment success. Interferons have been widely used in cancer; they pass BBB and act at JAK/STAT, PI3K/AKT/mTOR, and MAPKs cascades. We observed antiproliferative effects over GBM clones and cell lines in culture. U-87 MG is used as a model to understand the HeberFERON mechanism of action in GBM. We completed the first proteomic and label-free quantitative phosphoproteomic analysis after incubation of U-87 MG cell line with HeberFERON for 72h. The main contribution of this article is the description of phosphosites regulated in proteins participating in cell cycle, cytoskeleton organization, translation, autophagy, and DNA repair in a highly interconnected molecular network, where mTOR occupies a centric hub. Together with reported phosphosites, we described new ones and others with no associated kinases. Increased phosphorylation is mainly accounted by CDK and ERK kinases pointing to possibly new cascades regulations. This knowledge will contribute to the functional understanding of HeberFERON in GBM joined to general regulatory mechanisms in cancer cells.

## 1. Introduction

Interferons (IFNs) are cytokines with pleiotropic actions including antiviral and growth-inhibitory effects. These cytokines are the first line of defense against viral infections and have important roles in immunosurveillance for malignant cells [1]. Signaling crosstalk between IFN-α/β and -γ to induce stronger responses has been described [2]. IFNs type I and II transduce the signals after receptor binding through multiple cascades which includes: JAK/STAT, phosphoinositide 3-kinase (PI3K)/mTOR, and MAPKs [1, 3]. Much activation/deactivation of these cascades occurs through phosphorylation/dephosphorylation reactions. The last two cascades normally activate survival signals however in IFN response they function in a way that the outcome is antiproliferation, apoptosis, cell cycle arrest, or other processes leading to the decrease of cells growing and eventually their death [4]. Cells integrate information from multiple signaling pathways for a final proliferation outcome, so deciphering molecular mechanism is truly challenging.

A co-formulation of IFN-α2b and IFN-γ, in synergic proportions, known as HeberFERON has been approved for the treatment of skin basocellular carcinoma but also used off-label for other types of cancers including glioblastoma [5]. On the other side, due to its strong antiviral effect, HeberFERON has been more recently assayed in COVID-19 patients [6] and was included in the Cuban medical protocol for COVID-19 treatment [7].

Glioblastoma is the most lethal tumor of the brain with an average of 14 months of overall survival (OR) [8]. Different approaches have been assayed for Glioblastoma treatment with modest results in terms of OR, even though improvement in the quality of life of patients has been reported [9]. The use of HeberFERON in high-grade gliomas (1 anaplastic astrocytoma and 9 GBM) in a compassionate study prolonged the survival of patients to a mean of 34±14 months since diagnosis. The quality of life for all patients improved, following the Karnofsky Performance Scale (KPS) score and muscular power was improved in 50% of patients [10]. We have demonstrated an antiproliferative response of a panel of 34 glioblastoma-derived clones and several *in vitro* cell lines to HeberFERON [11]. Glioblastoma-derived cell line U-87 MG treated with HeberFERON has been chosen as a model.

Cell signaling is the biochemical process by which cells respond to perturbations in their environment, extracellular stimuli, or intracellular cues where the engagement of IFNs to their receptors would be one of those perturbations. Treatment of U-87 MG with HeberFERON causes an antiproliferative effect with a decrease of cell numbers in the treated culture. Cell signaling occurs through proteins post-translational modifications, including phosphorylation [12]. Protein phosphorylation at serine, threonine, and in <1% tyrosine, are one the most common post-translational modifications. Protein phosphorylation controls protein activity, localization, and protein complex formation [12, 13]. In humans, 518 protein kinases control phosphorylation. Dysregulation of kinase signaling pathways is commonly associated with various cancers [14]. Therefore along with changes in overall quantities of transcripts and proteins, the modification of phosphorylation sites is also an important way of regulation that should be taken into account in the biological effect of a drug.

Here we report the phosphoproteome and proteome changes of the U-87 MG cell line after the treatment with HeberFERON for 72h. Interpretation of these changes should derive into a better understanding of the molecular effects of HeberFERON over this glioblastoma-derived cell leading to an antiproliferation state.

## 2. Materials and Methods

### 2.1 Preparation of Samples for Proteomic experiment

U-87 MG cell line (ECACC 89081402) was grown in EMEM with 1.5 g/L of NaHCO_3_, 2mM of glutamine (Sigma, US), 50 μg/ mL of gentamicin sulfate (Sigma, US), and 10% of fetal bovine serum (Capricorn Scientific, Germany). Ten million cells per each condition were treated with HeberFERON (IFNα2b + IFNγ) at the concentration equivalent to IC50, for 72 hours. Three independent replicated cultures were treated or not (control) to experiment. After washing the pellet with PBS, cells were resuspended in 6M guanidine hydrochloride, 100 mM buffer Tris-HCl, 10 mM DTT in presence of proteases and phosphatases inhibitors. Cells were vortexed for 10 s and kept shaking for 15 min. Incubated for 2 hours at 37 ºC and later made up to 25 mM iodoacetamide and kept dark for 20 min at 25 ºC and centrifuged 3 hours at 60000xg. Proteins were processed by MED-FASP [15] with consecutive digestion using Lys-C and trypsin as described previously [16]. A small fraction of peptides from both digestions were desalted and later analyzed via LC-MS/MS for non-phosphopeptides identification and further proteome differential expression analysis. Phosphopeptides were enriched from each Lys-C and Trypsin derived digestions by using TiO_2_ beads with a 4:1 ratio (mg beads: mg peptides) as described previously [17].

### 2.2 LC-MS/MS and identification of peptides and phosphopeptides

Chromatographic runs for phosphopeptides and non-phosphopeptides were in home-made column (75 μm ID, 20 cm length), 120 min gradient, starting at 5 % B up to 30 % B in 95 min, then increase to 60 % B in 5 min, and up to 95 % in 5 min more. NanoLC was coupled to a Q-exactive HF mass spectrometer; mass range 300-1650 m/z was scanned using data-dependent acquisition. Each mass spectrum obtained at 60000 resolution (20 ms injection time) was followed by 15 MS/MS spectra (28 ms injection time) at 15000 resolution. Proteins and phosphopeptides were only considered when detected in at least two replicates in any of the groups.

Identification of peptides and proteins was based on the match-between-runs procedure by using Maxquant (v1.6.2.10) considering partial oxidation (M), deamidation (NQ), N-terminal acetylation (proteins), and phosphorylation (STY) as variable modifications. Alignment of chromatographic runs was allowed with default parameters (20 min time window and matching of 0.7 mins between runs). No fixed modifications were considered, as cysteines were not S-alkylated. Statistical analysis for the identification of phosphopeptides (class I sites with the probability of modification higher than 0.75) changing more than two-fold (p<0.05) were performed by Perseus (v1.6.2.2) after filtering for two valid values in at least one group [18]. Peptides with phosphorylation levels changing the same direction than at protein level were discarded.

### 2.3 Bioinformatic enrichment analysis

Enrichr [19] and Toppfun [20] were used for protein enrichment analysis with a p-value cutoff of 0.05. KEA2 [21], KSEA [22], and iPTMNet [23] were used for Kinase-Substrate Enrichment Analysis. KEA2 web tool (https://www.maayanlab.net/KEA2/) computes a Fisher Exact Test to distinguish significantly enriched kinases (p-values lower than 0.05), through statistical analysis. iPTMnet based on a set of curated databases like PhosphoSitePlus (http://www.phosphosite.org) and PhosphoEML (http://phospho.elm.eu.org), annotates experimentally observed post-translational modifications. KSEA app (https://casecpb.shinyapps.io/ksea/) based on a z-score transformation; assumes that the resulting scores were normally distributed. P-value is automatically calculated by the program. Sequence logos for those enriched motifs from phosphorylation sites with a p≤ 0.05 between control and experimental conditions were generated using WebLogo 3.7.4 (http://weblogo.threeplusone.com/). Phosphosite Motif analysis was also carried out using Phosphosite algorithm [24]. PhosphoSitePlus (http://www.phosphosite.org) was used for phosphosites additional information.

### 2.4 PPI analysis

All identified phosphoproteins were represented in a network context. STRING database (https://string-db.org/) (accessed 24 March 2021) was used to identify functional and physical associations between proteins. Only curated knowledge and experimental evidence were used as the source of interaction data in such analysis, and the confidence score was fixed at 0.4. The protein-protein interaction network was visualized using Cytoscape software (v.3.8.2); the MCL algorithm was used to identify clusters of tightly connected proteins within the network. Bisogenet (version 3.0.0) [25] was used to construct small networks with proteins participating in relevant biological processes.

## 3. Results

### 3.1 General Proteomic and Phosphoproteomic results

In this work, we studied the phosphorylation signaling involved in the antiproliferative effect of HeberFERON over the glioblastoma U-87 MG cell line. To survey proteome and phosphoproteome we treated U-87 MG with HeberFERON for 72 hours and enriched phosphopeptides from proteomes that were digested following the MED-FASP processing (Figure 1A). After phosphopeptides enrichment with TiO_2,_ we identified 23549 peptides (derived from 5692 proteins), 38 % (8900) of them were phosphopeptides; 6415 of these phosphopeptides were selected for statistical comparison among replicates. Considering a fold change higher than 2 for those Class I phosphopeptides (with a localization probability higher than 0.75), 412 phosphopeptides (p<0.05) and 143 phosphopeptides (p<0.01) were identified, corresponding to 359 proteins (Table 1S). A small fraction before phosphopeptide enrichment was separated for differential protein analysis. After LC-MS/MS 80764 peptides from around 7627 proteins were detected, and around 6000 of them were selected for statistical processing. With a fold change higher than 2, 333 proteins regulated by HeberFERON with p<0.05 were found; 122 were downregulated and 211 upregulated (Figure 1B, Table 2S).

**Fig 1.**
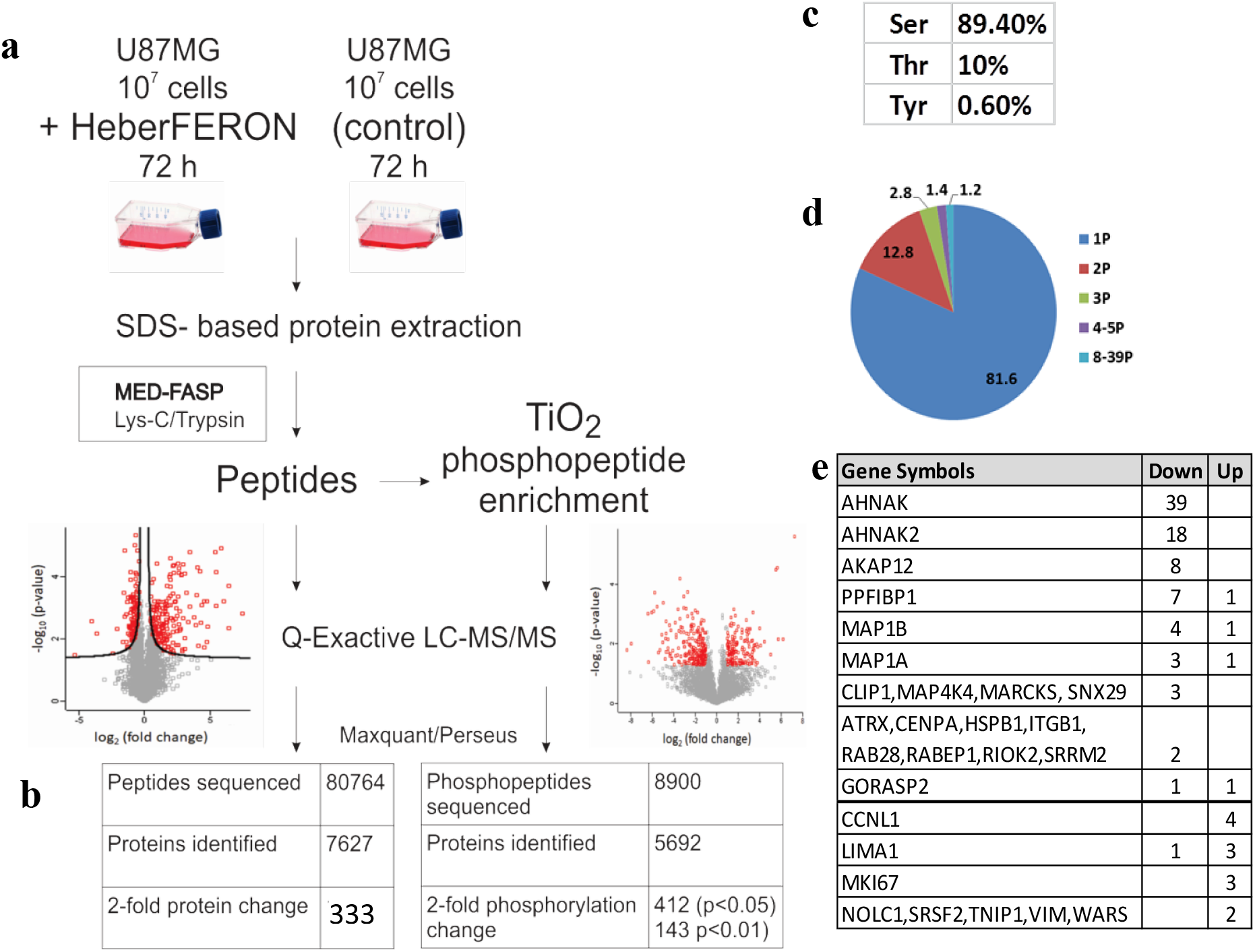
Experimental workflow and data processing for the identification of non-phosphopeptides and phosphopeptides regulated by HeberFERON. (**a**) Experimental workflow for sample processing, MS, and Data analysis (**b**) Comparison of identified non-phosphopeptides, phosphopeptides, and proteins (**c**) Distribution of phosphorylated amino acids in % (**d**) Number of phosphorylation sites per protein (**e**) Proteins with 2 or more phosphosites. Up and Down represent the partial increment or decrement of the phosphorylation to the non-modified peptide. Gene symbols are represented.

The distribution of pS, pT, and pY in phosphosites were 89.4, 10, and 0.6 %, respectively (Figure 1C). Peptides were predominantly singly and doubly phosphorylated, but some few proteins showed multi-phosphorylation of up to 39 sites (Figure 1D&1E). We identified 171 phosphosites increasing phosphorylation (from 141 proteins; 164 unique phosphosites) and 396 phosphosites decreasing phosphorylation (from 225 proteins; 344 unique phosphosites).

### 3.2 Enrichment Analysis

The enrichment analysis of proteome (Figure 2) with Enrichr tool highlighted biological processes (BP) such as IFNs signaling and cellular response to IFNs, negative regulation of viral life cycle or genome replication, and antigen processing and presentation via MHC. These processes are represented by classical encoding Interferon stimulated genes (ISG) like B2M, GBP2, IFIT1, IFITM3, ISG15, ISG20, IRF9, HLA-A, HLA-B, HLA-C, HLA-F, HLA-E, MX1, OAS1, OAS2, OAS3, OASL, SP100, and STAT1. The molecular functions (MF) of all proteins are mostly related to the double-stranded & single-stranded RNA binding (ex. RIG1 coding genes (DDX58), PKR (EIF2AK2)) and GDP binding, MHC class II receptor & aminopeptidase/peptidase/GTPase activities. In Cellular components stand out MHC class I and II complexes, protein complex components from the endoplasmic reticulum (ex.TAP1, TAPBP), Golgi, phagocytic vesicles, lysosome, and recycling endosome (ex.RAB8A, RAB35). Several proteins are shared among categories. Pathways enrichment by Toppfun also pointed to Interferon signaling pathways as the most statistically represented (Table 3S). Antigen processing and presentation, adaptive and innate immune systems, and the antiviral mechanism by IFN-stimulated genes are well represented. Additional pathways comprised the Signaling pathway from G-protein families (which includes kinases encoded by PRKCA and RPS6KA3); RAB GEFs exchange GTP for GDP on RABs membrane trafficking and Proteosome (more specifically Immunoproteasome components). Network analysis of the proteome data showed highly interconnected biomarkers from STAT1/STAT2 to antiviral (Figure 1S-a) and MHC processing/presentation and immunoproteasome (Figure 1S-b) components.

**Fig 2.**
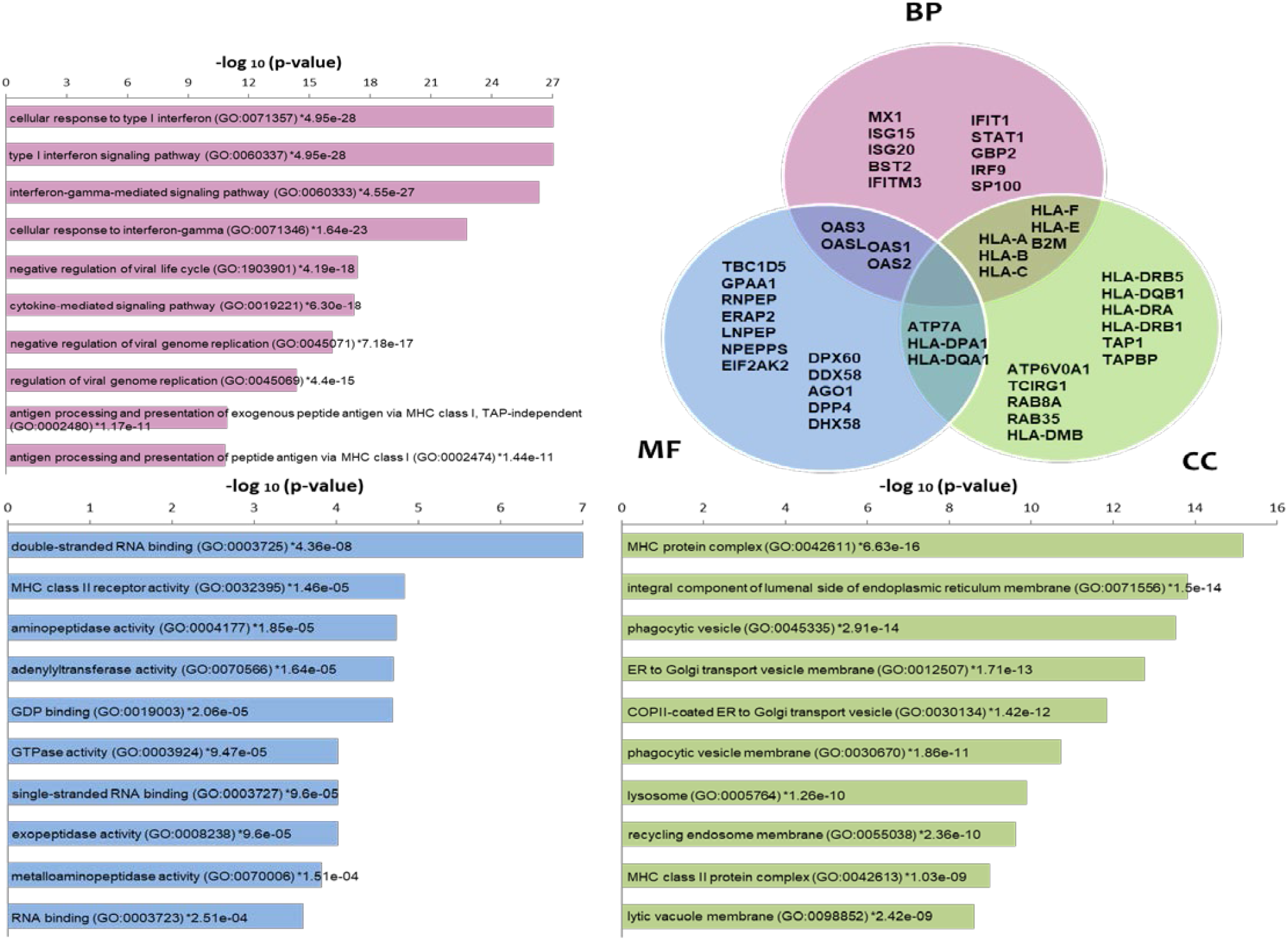
Enrichment analysis of proteomic data using Enrichr tool. Main processes are shown in the categories Biological Processes (BP), Molecular functions (MF), and Cellular components (CC) as bars. A Venn diagram is also used to signify some of the proteins included in each category as well as some that are common among categories. Gene symbols are used in the Venn diagram.

Phosphoproteins enrichment analysis using the Toppfun tool detected relevant biological processes (Figure 3A). Among the most represented we found the cytoskeleton organization followed by chromosome organization and cell cycle while others such as autophagy, RNA splicing, regulation of GTPase activity and regulation of apoptotic signaling pathway, translation, DNA replication, DNA repair, small GTPase mediated signal transduction, and response to unfolded protein are shown with different levels of importance according to their – log (p-value) (Figure 3A). Looking at the pathways using the same tool, Cell Cycle (more specifically Mitotic) is among the most relevant paths, as well as, those related to Gene Expression, rRNA processing, apoptosis, and DNA repair (Figure 3B). mTOR signaling pathway seems to be regulated by HeberFERON changing phosphorylation levels of different phosphoproteins from this pathway.

**Fig 3.**
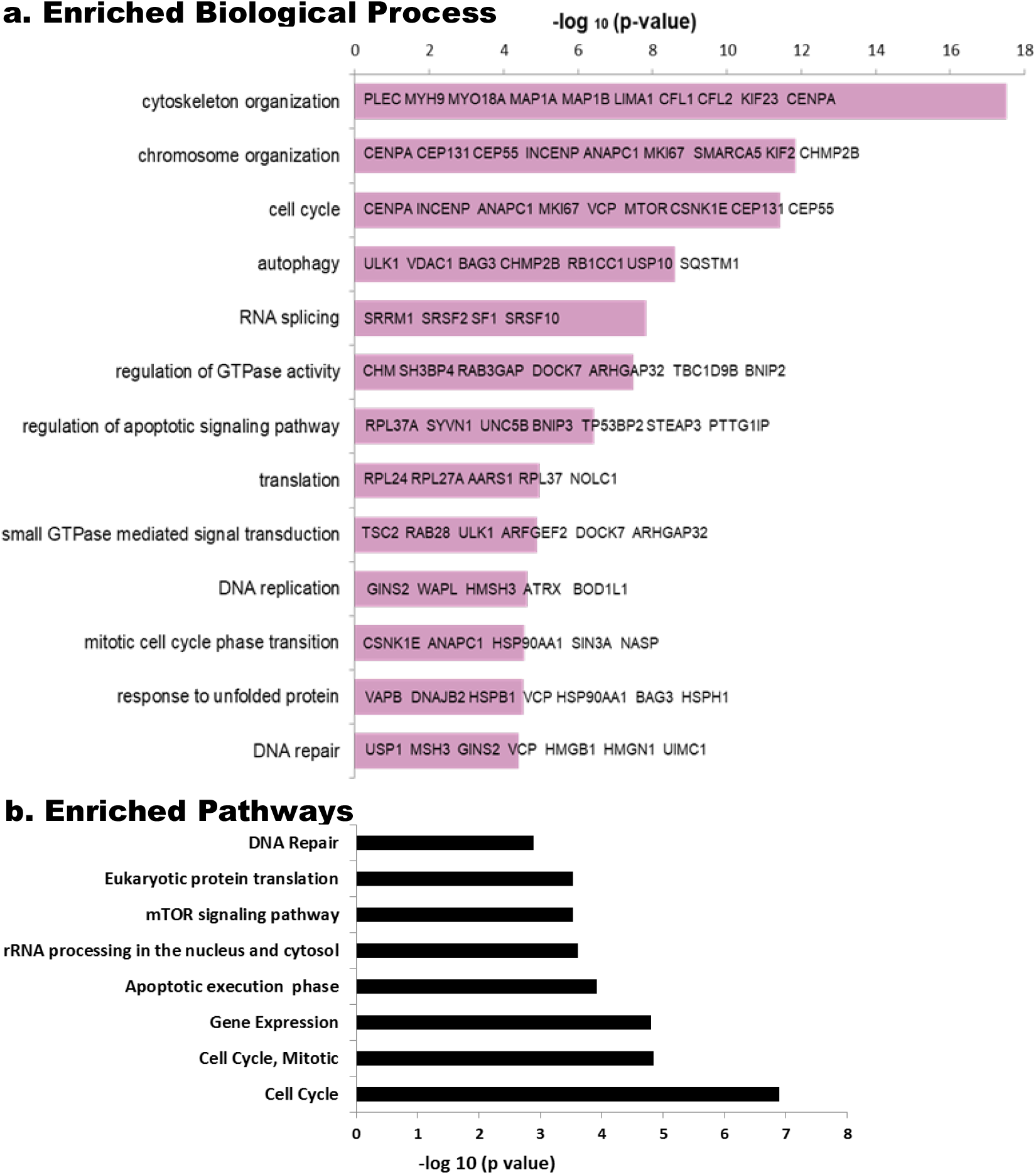
Enriched Biological processes and Pathways in phosphoproteomic data using Toppfun tool. The reciprocal of Log (p) is tabulated for each Biological Process (a) and Pathways (b). Some of the principal components of the Biological Process, as gene symbols, are shown in bars.

### 3.3 Analysis of significant kinases through regulated phosphosites

Figure 4 shows sequence motifs with a LogoSequence graph for phosphosites that increased (UP) or decreased (DOWN) phosphorylation with HeberFERON (Figure 4A). Analysis of these motifs was carried out using the Motif-All algorithm in PhosphoSite Plus (Table 4S). Motifs with the highest probability for phosphosites increasing phosphorylation looks like X-**S-P**-X-K. This Proline-directed phosphosite is typical for CDK and ERK proteins. To a lesser extent, mTOR motif like X-**T**-**P**-X-X-S-K was also predicted. KSEA program (Figure 4B) similarly pointed to CDK1, CDK2, MAPK3/K1 (alias ERK1/2), CDK3/4/6/7, as those kinases with the highest z-scores based on the collectively increased phosphorylation status of their phosphosites with a p< 0.05. This program showed other kinases as AURKA, RAF1, and ROCK1. On the other hand, motifs with the highest probability for phosphosites decreasing phosphorylation are more variable but we observed a general motif as RK-RK-K-**S**-FLV-KR/X, corresponding to protein kinases (PK) ABC family. KSEA showed kinases with the lowest z-Scores based on the decreased phosphorylation status of their phosphosites including PRKCA (PKA) and PRKCB (PKB). Additional kinases with z-scores from 0 to |1| could be also responsible for phosphosites modification. These are the cases of NEK2, RPS6KA2, and GSK3B regulating phosphosites with phosphorylation increases and AKT1, PKD1, AURKB, EIF2AK2, PRKCE, PRKDC, PLK1, EEF2K, RPS6KA1/A3 for phosphorylation decreases.

**Fig 4.**
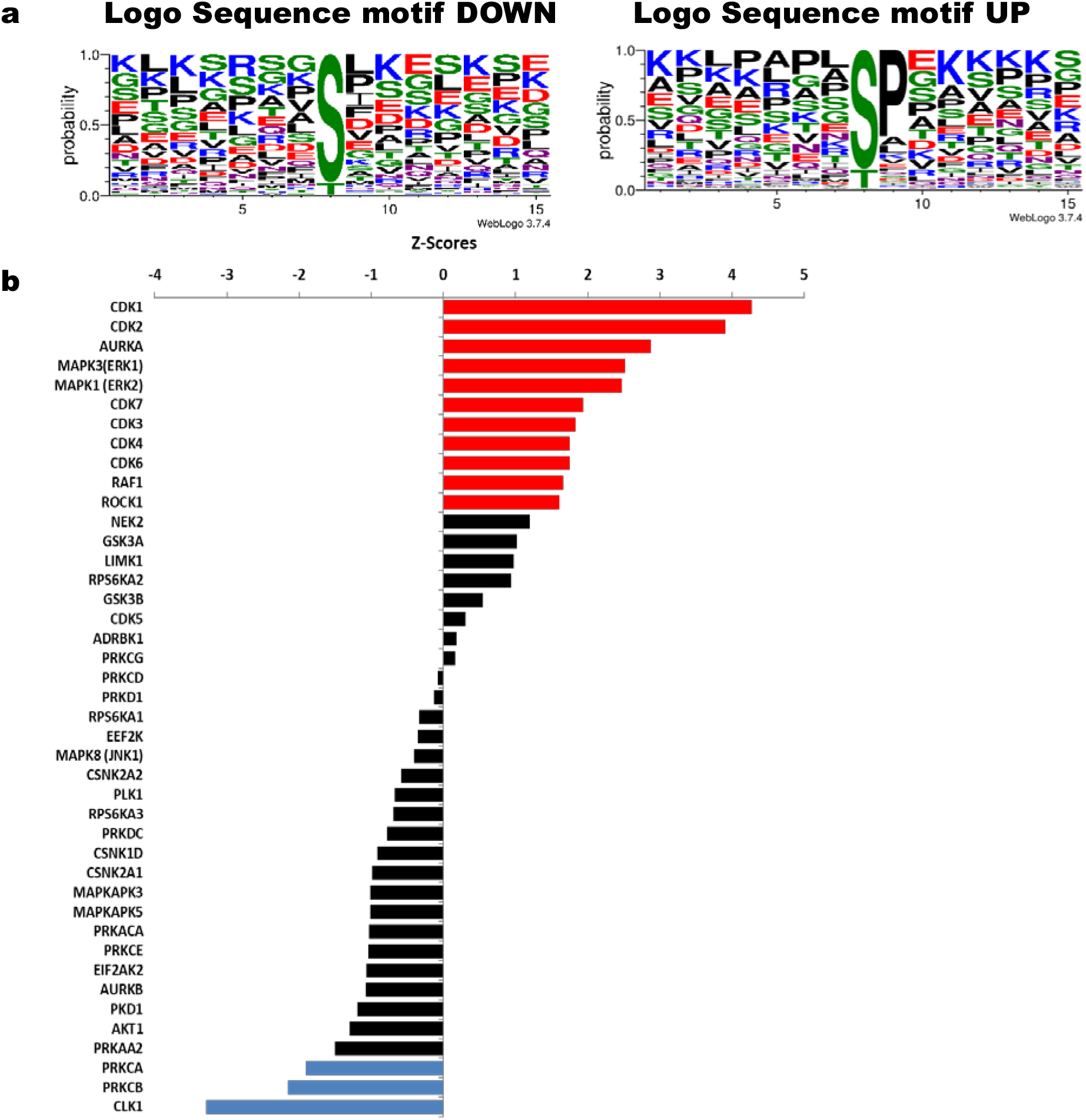
Sequence motifs logo and kinases analysis from phosphosites with increased or decreased phosphorylation after U-87 MG treatment with HeberFERON. **(a)** Logo Sequence motifs found for phosphosites that increased (UP) and decreased (DOWN) phosphorylation, (**b**) Analysis of kinases using KSEA tool taking into account phosphosites statistically regulated; red (for sites with increased phosphorylation) and blue (for sites with decreased phosphorylation) for kinases with Z scores with statistically significance; in black other kinases with less significance. Gene symbols are displayed.

Table 1 listed some of the most interesting phosphosites on substrates regulated by enriched kinases from KSEA analysis. Several phosphosites were obtained after the analysis with KEA2 and/or iPTMNet programs. Following kinase predictions, most of the phosphosites changing phosphorylation levels were CDK1/CDK2/PRKCA/PRKCB substrates, and some of them were also regulated by ERK1 & ERK2, AURKA, or CDK3/4/6/7. We found regulated phosphosites in proteins with important roles in cell cycle and proliferation (i.e. RB1_S807, MKI67_S584, INCENP_S421, CENPA_S19, MTOR_S1261), cytoskeleton organization (i.e. MAP1B_T1788, MARCKS_S27, MARCKS_S29), translation (EIF5B_S222, RPL12_S38), DNA repair, and autophagy (i.e. BRCA1_S114, RAD9A_S328, RFC1_T506, SQSTM1_S272, MTOR_S1261) and in multifunctional proteins like NPM1/B23, HSPB1/HSP27, AKAP12, and AHNAK. Substrates of RAF1 & ROCK1 (VIM_S420 and CFL1_S3) and GSK3B (several sites on MAP1B, CLIP1/2) were regulated. PLK1, AURKB, AKT1, EIF2AK2 (PKR), RPS6KA3, PKD1, and other PKC kinases are related to the decreased phosphorylation in phosphosites on HSPB1/HSP27, mTOR, and NPM1/B23.

**Table 1.**
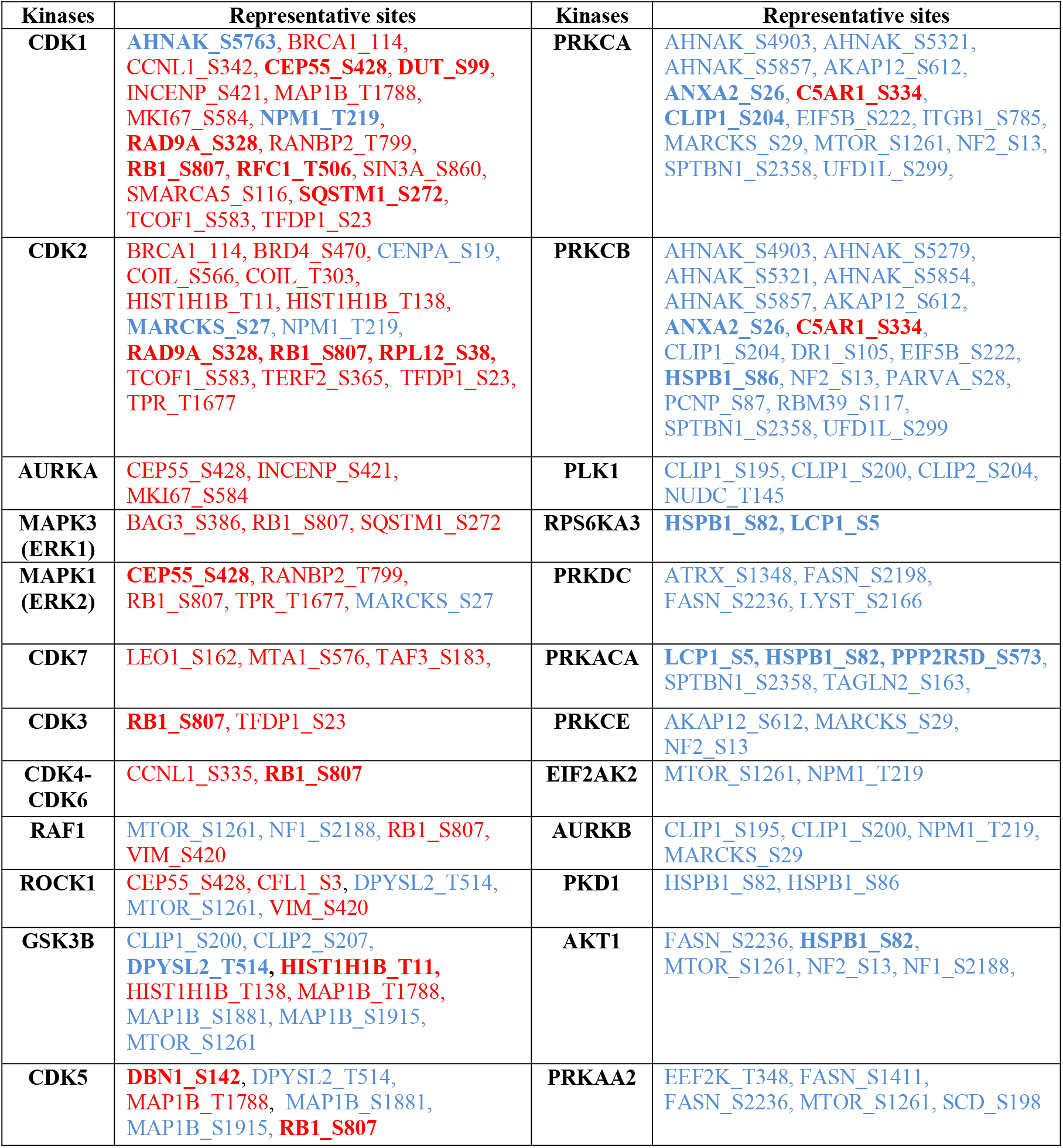
Principal substrates and phosphosites regulated by kinases following KSEA analysis. Phosphosites emphasized in bold letters were also obtained from KEA2 and/or iPTMNet analysis. Phosphosites increasing phosphorylation are in red and decreasing phosphorylation in blue. Gene symbols and the position of phosphosite in each protein are shown.

There are numerous regulated phosphosites that have not been described or that have been described with no associated kinases for phosphorylation. Table 2 shows some of those sites obtained after iPTMNet analysis, divided by Biological Pathways. Several of these phosphosites are on proteins participating in Cytoskeleton organization and Cell Cycle (i.e MAP1A and MAP1B, MYH9, MYO18A, and 9B, LIMA1, MARCKS, KIF23, and PCM1), Autophagy (i.e RAB7A and RB1CC1), Translation (i.e EIF4G1 & G3), RNA Splicing and on multifunctional proteins like AHNAK, AHNAK2, and AKAP12, which show several phosphosites decreasing phosphorylation. We found a decreased phosphorylation in the mTOR signaling member, GSK3B_S25, and an increased phosphorylation in the new phosphosite IFNAR1_T328.

**Table 2.**
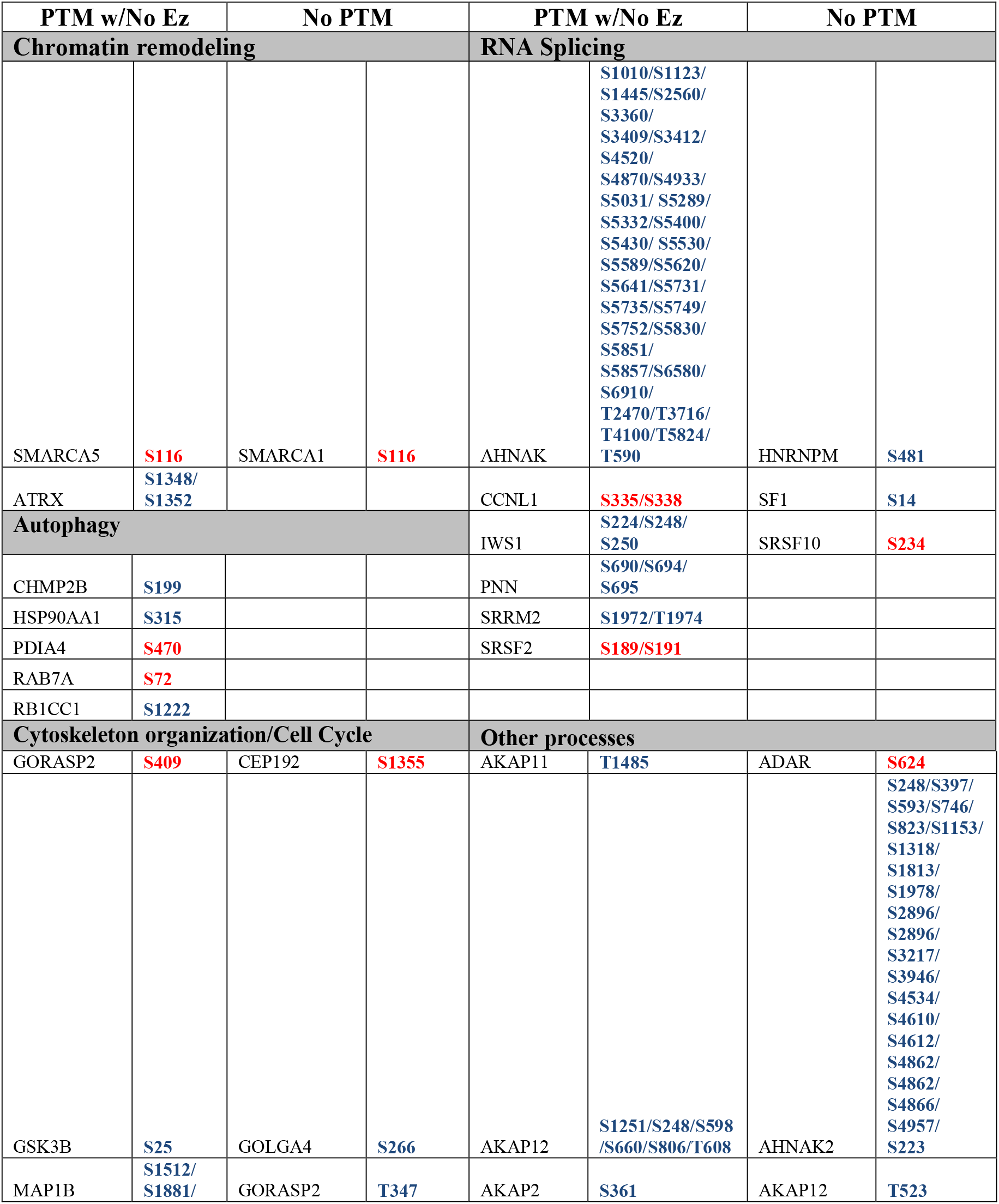
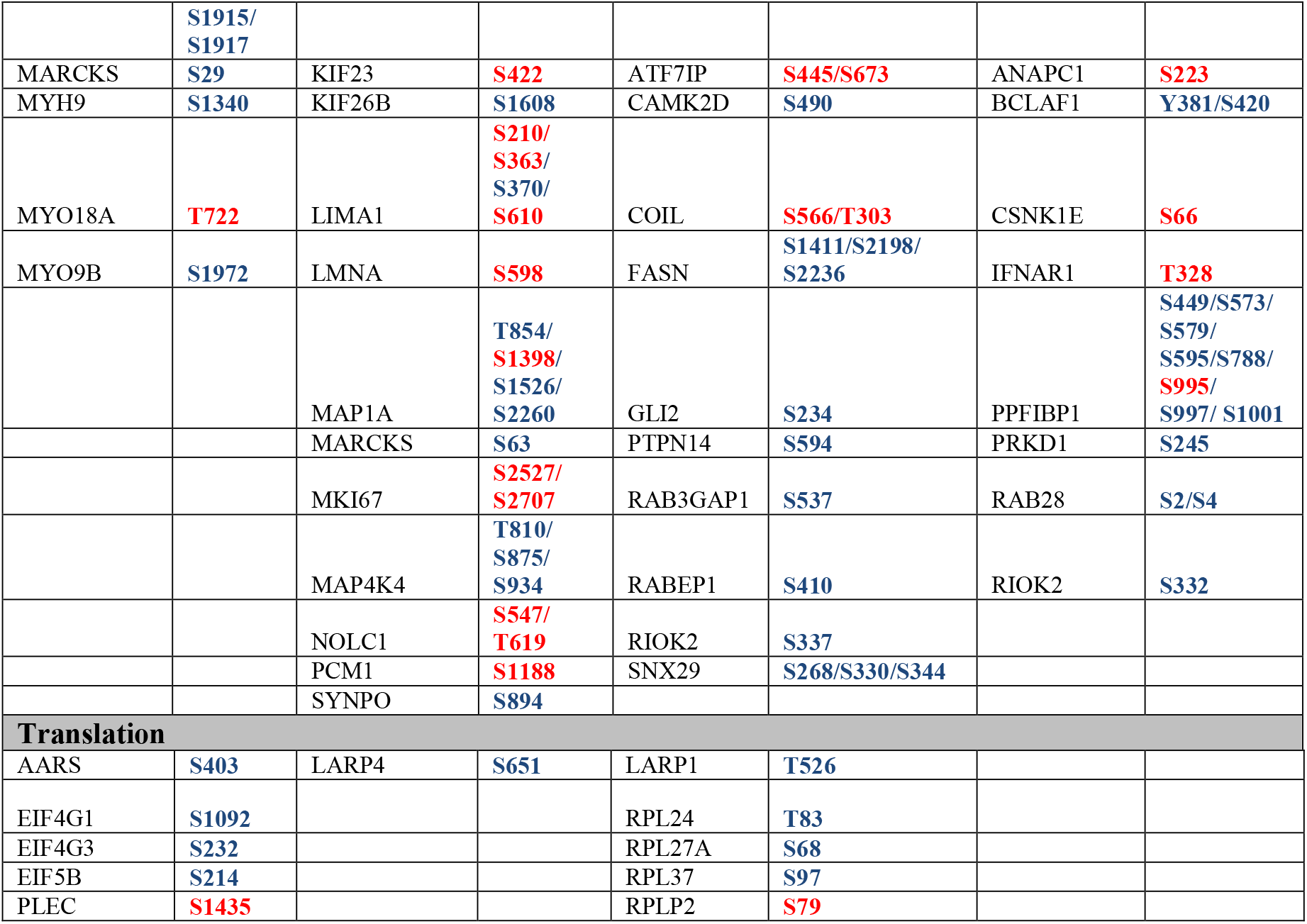
Substrates and regulated phosphosites not described before (No PTM) or described with no kinase-associated (PTM w/No Ez) following iPTMNet analysis. Phosphosites with increased phosphorylation are in red and with decreased phosphorylation in blue. Gene symbols and the position of phosphosite in the proteins are shown. They were grouped by biological process.

### 3.4 Network analysis

The network interconnections among proteins with phosphosites regulated by HeberFERON were analyzed using STRING on the Cytoscape framework (Figure 5). For a better understanding, we grouped proteins by Biological Processes according to STRING classification. There are nodes with high degrees of more than 15 connections with other proteins, such as NPM1/B23 and CENPA in Chromating remodeling; CENPA as part of Cell Cycle too, proteins MYH9, MYO18A, BRCA1 or HSPB1/HSP27 in the cytoskeleton organization and VCP & HSP90AA1 acting at autophagy. In RNA splicing, translation, and autophagy processes, there are highly connected clusters. Translation show several 60S ribosomal subunit and tRNA ligases proteins with distinct regulations, as well as, components of the initiation complex (EIF4G1_S1092, EIF4G3_S232) or the elongation complex related to MTOR-RPS6KA3 pathway (EEF2_T57, EEF2K_T348) decreasing phosphorylation. RPS6KA3 protein was downregulated and phosphosites controlled by this kinase (i.e HSPB1_S82 and LCP1_S5) congruently decreased phosphorylation. MTOR occupies a centric position interacting with several other proteins in processes such as autophagy (TSC2, RB1CC1, HSP90AA1, ULK1, and SQSTM1), cell proliferation (GSK3B, BRCA1), or translation (*not shown*). AHNAK, with 39 phosphosites decreasing phosphorylation, is located in the frontier between RNA Splicing (interacting with SF1) and cytoskeleton organization (interacting with Periplakin (PPL), Golgi protein GORASP2, and the Ephrin type-B receptor 2 (EPHB2)). Interactions with MYH9, LIMA1, HSPB1/HSP27, EEF2, and ANNXA2, could also occur although not shown in this network. Other proteins with multiply regulated phosphosites (i.e AHNAK2, ANAP12, Leprin beta 1 (PPFIB1), SNX29) are not connected in the main interaction network but most of the proteins, even those with no previous report of phosphosites or with no clear kinase association, are included.

**Fig 5.**
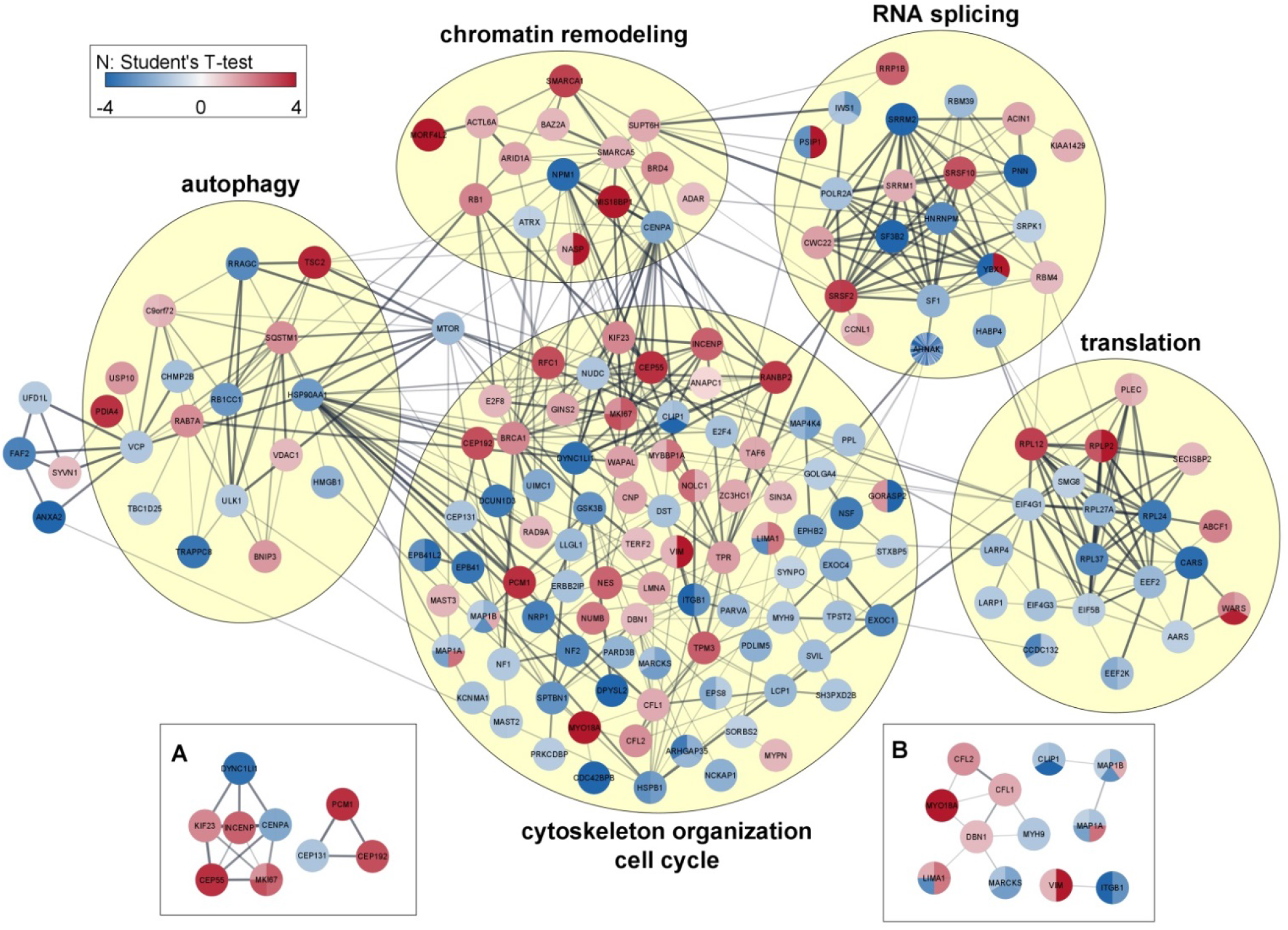
Network association among proteins with regulated phosphosites. The relations are grouped by Biological processes. For each protein, the color indicates the direction of regulation for up or down and the relative value of that change as it is indicated with the color intensity bar. Clusters of interest are presented in **a**: cell cycle significant proteins and **b**: cytoskeleton significant proteins. Gene symbols are shown.

## 4. Discussion

In this article, we present the first label-free quantitative proteomic and phosphoproteomic results after the treatment of the glioblastoma-derived cell line U-87 MG with HeberFERON for 72 hours.

IFNs signaling start with the binding of the IFNα and IFNγ to their receptors and follow the canonical JAK/STAT pathway, where STAT1 and STAT2 in association with IRF9 initiate the signal transduction cascade and different complexes translocate to the nucleus to activate the expression of genes encoding proteins involved in diverse processes. STAT1, STAT2, and IRF9 increased protein expressions with HeberFERON. Many interferon-stimulated genes from ISRE or GAS promoter sites can explain direct antiviral, innate immune, and antiproliferative responses [26-28]. The increase of several classes I and II MHC molecules, proteins participating in MHC-peptide presentation, post-translational modifications, or in proteasomes explains the activation of adaptive immunity [29]. Activation of IFIT3 [30], 2’-5’ OAS and RNase L [31], PKR [32], and IRF9 [33] could contribute to the antiproliferative effect over U-87 MG cells. In glioblastoma-derived T98G cells, IRF9 overexpression enhanced the antiproliferative activity of IFN-α2c in resistant cells and induced apoptosis [33].

HeberFERON exerts interferons synergic and longer time effects over U-87 MG [5]. This could be explained by the crosstalk between both IFN cascade signaling [2] and the IFN signaling feedback controls including receptor recycling [34]. This experiment gives new clues. Upon IFNα stimulation, PKD2 is recruited to IFNAR1, phosphorylated by TYK2 which in turn phosphorylates IFNAR1 on S535 and S539 residues. These phosphorylations are important in the internalization of the IFNAR complex. After internalization and arrival in early endosomes, degradation in lysosomes is used for IFNAR1 to control signaling. Here, we found the site not previously described IFNAR1_S426 with increased phosphorylation. Nevertheless, it is suggested IFNAR2 is recycled back to the plasma membrane by a retromer complex [34]. This complex is assembled by a first trimer sub-complex, made of vacuolar proteins sorting-associated protein 35 (VPS35)-VPS29-VPS26, that binds the early endosome phosphoinositide PI(3)P, through a domain present in sorting nexin (SNX), and together with the small GTPase RAB7A, assembles the second sub-complex of the retromer. IFNAR2 can interact with RAB35 and VPS26A, VPS29 and VPS35. In our data VPS26B and RAB35 decreased and increased protein expressions, respectively. Phosphosites VPS26A_S2 and S268/S330/S344 on SNX29 decreased phosphorylation; RAB7A_S72 increased phosphorylation. Only the phosphorylation in RAB7A has been associated with LRRK1 to regulate cargo-specific trafficking [35]. Controlling the residency time of internalized IFNAR complex in the endosome, the retromer is directly implicated in the fine tuning of JAK-STAT signaling duration and downstream transcription outputs thus the long-term effects of HeberFERON could be explained by the persistence of receptor-bound IFN-α2 inside endosomes and IFN signaling from this compartment for days even when the negative regulators ISG15 or USP18 were missing [28]. The regulation of components contributing to IFNAR2 recycling and the new phosphosite on IFNAR1 could explain the higher and longer effects of HeberFERON compared to IFNs type I or II independently [5].

IFNs could also achieve an antiproliferative response through non-canonical signaling including PI3K/AKT/mTOR and MAPK cascades. ERK1 and ERK2, from the MAPK protein family, can impact cell cycle progression, cytoskeleton organization, and mRNA translation [36]. mTOR impacts translation, cytoskeleton, and autophagy [37]. Protein kinases (PK) ABC family also participate in these signaling leading to proliferation control [38].

Changes in phosphorylation of key proteins in the cell cycle support this is a very important biological process targeted by HeberFERON [39]. The histone H3 variant centromere protein A (**CENPA**) specified accurate segregation of sister chromatids during mitosis [40]. Phosphorylation of CENPA on S17 and S19 by CDK2 is important for chromosome segregation and unphosphorylated variants result in mitotic errors [40]. HeberFERON diminished phosphorylation of both sites (S17 and S19) on CENPA, a protein highly connected in the phosphoproteomic network. **INCENP** ensures the correct chromosome alignment and segregation, as a scaffold protein regulating the chromosomal passenger complex localization and it is required for chromatin-induced microtubule stabilization and spindle assembly [41]. INCENP_S420 and INCENP_S424 increase phosphorylation in mitosis [42], but HeberFERON increased phosphorylation of INCENP_S421. This site has been described in Phosphosite Plus phosphorylated by CDK1 with no biological function. Increased phosphorylation of phosphosites unreported in the APC/C component **ANAPC1_S223** and the kinesin **KIF23_S422** [41] occurred with HeberFERON. The decreased of CENPA, INCENP, APC/C-CDC20 and KIF23 gene expressions with a significantly fold change respect to individual IFNs was observed in a microarray experiment, reinforcing their roles in the G2/M arrest observed [39]. Three phosphosites (S584, S2527, and S2707) with no biological function described in **MKI67**, increased phosphorylation [43]; only S584 is predicted to be a CDK1 substrate by NetworKIN. This protein is required to maintain mitotic chromosomes dispersed thus it is present at the highest level in the G2 phase and during mitosis. Phosphorylation at both S807 and S811 in **RB1** by CDK3/cyclin-C promotes G0-G1 transition [44]. HeberFERON treatment only increased phosphorylation of the first one.

The Ser/Thr kinase mTOR cascade is commonly upregulated in cancer due to loss of the tumor suppressor PTEN, as it is in U-87 MG [45]. In tumor cells, PI3K/mTOR axis is necessary for the induction of apoptosis after treatment with IFN-α [4]. In our data, **mTOR** occupies a centric position in the phosphoproteomic network. PI3K/AKT as well as the mitogen-activated protein kinase (MAPK) pathways engages mTOR to regulate a variety of components of the translational machinery, where mTORC1 regulates mRNA translation by phosphorylation of components of the initiation translation complex. Here, **EIF4G1_S1092** and **EIF4G3_S232** decreased their phosphorylations whose functional consequences are unknown, but the translatome stoichiometry changes could be a mechanism derived from mTOR influence [46]. In this same signaling branch, the ribosomal protein S6 kinase alpha-3 (gene **RPS6KA3**), a member of the RSK family that regulates translation through RPS6 and EIF4B phosphorylation, was found with decreased protein expression. TSC/Rheb signaling promotes mTOR S1261 phosphorylation, then promotes mTORC1-mediated autophosphorylation and substrate phosphorylation to stimulate cell growth [47, 48]. Unreported site **TSC2_S1743** increased phosphorylation and the phosphorylation of mTOR_S1261, performed by PKC alpha, GSK3B, RAF1, ROCK1, or AKT1, is decreased by HeberFERON. Thus, we should expect the activity of downstream substrates of mTOR, depending on S1261 phosphorylation, to be affected. **EEF2K_T349** and **EEF2_T57** diminished their phosphorylation. Calmodulin-dependent eukaryotic elongation factor 2 kinase (eEF-2K) _T349 autophosphorylation appears to control the catalytic output of active eEF-2K, contributing more than 5-folds to its ability to promote eEF-2_T57 phosphorylation, impeding protein synthesis. How this could be a rescue response due to the effect of HeberFERON over cell proliferation should be further investigated.

The cytoskeleton is a complex network of microtubules, actin, and intermediate filament that contributes to cell growth and migration. mTORC2 controls the actin cytoskeleton. mTORC2-associated interactome showed actin-binding proteins and microtubule-associated proteins significantly changed depending on mTORC2 activity level in glioblastoma including Myosin-9/MYH9, Plectin/PLEC, and MAP1B [49]. Microtubules are composed of alpha- and beta-tubulin heterodimers that assemble into protofilaments with numerous microtubule-associated proteins (MAPs) that bind along with the microtubule matrix [50]. MAP1 family associates with both microtubules and actin and their phosphorylation regulate the filamentous cross-bridging between microtubules and other skeletal elements. **MAP1A** showed three sites that diminished (T854/S1526/S2260) and one that increased (S1398) phosphorylation. **MAP1B** also showed four sites with decreased (S1512/S1881/S1915/S1917) and one with increased (MAP1B_T1788) phosphorylation, all of them not described before or with no biological function associated to those changes in PhosphoSite Plus. Deficiencies in expression or lack of phosphorylation of MAP1B by GSK3β led to abnormalities during brain development. MAP1B_T1788, MAP1B_S1881, MAP1B_S1915 are predicted substrates for GSK3B but also for CDK1, CDK5, GSK3A and other kinases in NetworKIN. Additional **GSK3β** regulated sites on cytoskeleton proteins (CLIP1_S200, and CLIP2_S207) diminished phosphorylation. The proline-directed protein kinase GSK3β [51] looks like another important hub in the HeberFERON action and it is functionally related to mTOR signaling [52]. This kinase is inhibited by phosphorylation of GSK3β_S9 but instead, we found a decreased phosphorylation on GSK3β_S25 with unknown biological relevance or responsible kinases [51]. Another member of microtubule-associated proteins, **MAP4**, decreased protein expression. Additional actin-binding proteins were regulated by phosphorylation after HeberFERON treatment including **MARCKS, CFL 1** and **2, Eplin (LIMA1), MYH9**, and **MYO18A**; most of them highly connected in the phosphoproteome network. Phosphosites changing phosphorylation in Myosin 9/MYH9 and MYO18A have unknown effects [53]. Eplin coordinates actin and myosin dynamics throughout cell division by increasing the number and size of actin stress fibers, it is phosphorylated at the C-terminal region by ERK1/2 which reduces its association with F-actin and contributes to actin filament reorganization and cell motility enhancing [54]. After HeberFERON treatment, three sites increased phosphorylation in this protein (S210, S363, and S610) and one decreased (S370) but only S362 is reported phosphorylated by ERK1 with a downstream effect related to cytoskeleton organization and cell motility. It has been reported compensatory and synergistic effect of double phosphorylation at Eplin_S362 and S604 residues and it is potentially possible that other residues may also be involved in the regulation of Eplin [54]. HeberFERON increased the phosphorylation of S3 in CFL1 and CFL2 [55], which inactivate these proteins in their role over the normal progress of mitosis and cytokinesis. MARCKS is the most prominent substrate for PKC [56]; all these sites decreased phosphorylation: MARCKS_S27 regulated by CDK2 and ERK2, MARCKS_S29 regulated by PRKCG, PRKCD, and other kinases, and the unreported MARCKS_S63. Finally, class-III intermediate filament protein Vimentins (**VIM**) and the small heat shock protein HSP27 (**HSPB1**) are also highly connected in the network. In U251 glioblastoma-derived cell line interaction between VIM and GSK3β was demonstrated with implications in cell migration [57]. VIM_S339 and VIM_S420 increased phosphorylation, but both sites have no downstream biological function reported. NetworKIN predicts phosphorylation of S420 by RAF1 and ROCK1. HSP27 plays a role in stress resistance and actin organization functioning as a molecular chaperone and maintaining denatured proteins in a folding-competent state [58]. Phosphorylated HSP27 impairs its ability to protect against oxidative stress and HeberFERON decreased phosphorylation in S82 and S86. S82 is phosphorylated *in vitro or in vivo* by AKT1, MAPKAPK2 and P70S6KB with a biological function related to cell growth and cytoskeletal reorganization but phosphorylation at S86 has no known biological function associated. Cytoskeleton phosphoproteins showed several interactions among them and with proteins participating in cell cycle, translation, autophagy, DNA repair, chromatin organization, or splicing which could contribute to limit cellular proliferation.

GSK3β and mTORC1 are also critical regulators of autophagy downstream of the PI3K/AKT pathway [48]. Autophagy is a crucial lysosomal-dependent cellular degradation process for cell survival under extreme conditions, preserving cells from further damages [59]. But, autophagy also plays a death-promoting role, as a tumor suppressor mechanism in cancer and together with apoptosis could contribute to limit cancer cell proliferation. In response to starvation or after a pharmacological blockade, the mTORC1-dependent phosphorylation sites in ULK1/2 are rapidly dephosphorylated, which stimulates ULK1/2 autophosphorylation and phosphorylation of both Atg13 and RB1CC1 leading to autophagy induction. In this process, the HSP90-CDC37 chaperone complex selectively stabilizes and activates ULK1. Phosphorylation sites on the highly connected HSP90A (HSP90AA1) regulate the mechanism of HSP90 or affect the interaction between HSP90 and its co-chaperones and clients, many of which are protein kinases [60]. After HeberFERON treatment, phosphosites unreported or with unknown biological functions in **RB1CC1** (S516), **ULK1** (S623), and **HSP90A** (S315) decreased phosphorylation. Selective autophagy, in cooperation with the ubiquitin-proteasome system, acts as an ON-OFF switch of cell cycle progression via controlling the stability of several regulators, thus cell cycle proteins such as CDKs, CKIs, and checkpoints could regulate autophagy and vice versa [61]. **SQSTM1** is one of the receptors participating in selective macroautophagy recognizing CDK1, cyclin D1, p27, among others, and recruiting them into autophagosomes. Their relation with CDK1-Cyclin B1 has positioned SQSTM1 as a node for the control of cell survival and cell transit through mitosis [62]. Here, SQSTM1_S272, phosphorylated by CDK1 and MAPK P38 Delta, increased its phosphorylation. The ATPase enzyme **VCP** is also involved in the maturation of ubiquitin-containing autophagosomes and the clearance of ubiquitinated protein by autophagy and shows several network connections. As part of the complex containing NPLOC4-UFD1-VCP binds ubiquitinated proteins and export misfolded proteins from the endoplasmic reticulum to the cytoplasm, where they are degraded by the proteasome. It acts as a negative regulator of type I interferon production by the interaction of the NPLOC4-UFD1-VCP complex to **RIG-I/DDX58**. Disruption of the complex enhances RIG-I antiviral signaling [63]. Proteomic results showed the increase of RIG-I/DDX58 with HeberFERON and the phosphosite VCP_S13 decreased phosphorylation. Additional elements supporting the role of autophagy in the HeberFERON action come from phosphorylation changes in centrosome proteins: **PCM1_S1188, CEP131_S47, CEP55_S428, and CEP192_S1355**. PCM1 acts as an autophagy receptor participating in the autophagosome formation and leading to regulatory feedback pathways [64-66] and the relation among centrosome, autophagy, and cell cycle is reported [66]. Macroautophagy is indeed involved in the replication stress response and DNA repair pathways [67]. The histone chaperone **NPM1** with multiple roles [68] has a high degree of connection in the phosphoproteome network. NPM1_T219 diminished phosphorylation after HeberFERON treatment. This site together with T199, T234 and T237 is phosphorylated before mitosis by the CDK1-cyclin B complex and lack of phosphorylation or NPM expression result in spindle checkpoint activation due to severe mitotic defects [69]. Furthermore, other phosphoproteins controlled by CDK1 and CDK2 phosphorylation and participating in the DNA repair process [**BRCA1, UIMC1, RAD9A**, and **RFC1]** modified phosphosite phosphorylation status after HeberFERON treatment. The highly interconnected E3 ubiquitin-protein ligase BRCA1 showed increased phosphorylation in BRCA1_S114 which is important for fork and nascent DNA strand protection [70].

Lastly, we found proteins with scaffolding properties that showed several sites with decreased phosphorylation, including **AHNAK** (39 sites), **AHNAK2** (18 sites), **AKAP12** (8 sites), and **Liprin beta1** (PPFIBP1, 7 sites). AHNAK could be a regulator of PKC-alpha activity [71] and, in confluent cells, it is phosphorylated by protein kinase B (PKB)/AKT. This results in the translocation of AHNAK to the plasma membrane where it forms a protein complex with Annexin 2 (*ANXA2*)/S100A10 and the actin cytoskeleton contributing to cell-cell contacts. HeberFERON decreased phosphorylation of ANXA2_S26, which is related to actin remodeling. Treatment with PI3K inhibitor, upstream of PKB/AKT, resulted in decreased levels of phosphorylated AHNAK located in the nucleus. Decreased phosphorylation of a few sites (S4903, S5279, S5321, S5854, and S5857) is possibly modified by PKC kinases while AHNAK_S5763 is predicted to be regulated by CDK1. The future implication of this protein in cell growth through interferon non-canonical signaling should be studied.

U-87 MG as a tumoral cell promotes the proliferation through different signaling pathways (MAPKs, PI3K/Akt/mTOR, PKA, and PKC) where crosstalk between signaling pathways is common in cell regulation. HeberFERON acts over most of these signaling and fine-tunes the biological responses to achieve an antiproliferative effect at 72h of treatment. Most of the substrates phosphorylation increases were provoked by CDK1/CDK2 and ERK1/2, kinases with a recognized role on cell cycle progression and survival pathways. These two cascades control cell proliferation under very strict regulation including positive and negative feedback loops and reciprocal regulation which promote oscillations and dynamism necessary to adjust regulation [72, 73]. This fine-tuning could lead cancer cells to arrest or rescue outcomes, using mechanisms as autophagy or DNA repair. The balance through an arresting response can be a mechanism HeberFERON uses.

In this paper, for the first time we obtained answers about the main mechanism used by HeberFERON in U-87 MG to achieve an antiproliferative effect, through the proteomic and phosphoproteomic data (Figure 2S). Together with antiviral mechanisms, IFNα/γ co-formulation impact over the cell cycle, cytoskeleton organization, translation and RNA processing, autophagy, and DNA repair using several signaling pathways and highly interconnected phosphoprotein hubs, where mTOR occupies a centric position impacting all the mentioned biological processes. IFN canonical and non-canonical pathways converge to kinases as CDK1/CDK2, ERK1/2, PK ABC, mTOR, and GSK3 that regulate an enormous quantity of sites, most of them either not been described before or associated with biological functions. Furthermore, some of the regulated phosphosites could contradict the antiproliferative effect, suggesting a possible rescue response at 72h. The study of the phosphorylation changes will contribute to our future understanding of how HeberFERON succeeds with cellular proliferation arrest through novel proteins functions and interconnections in glioblastoma cultures. Future studies of the main biomarkers and pathways at earlier and later time courses could deeper clarify the mechanisms.

## 5. Acknowledgments

The authors are grateful to Prof. Matthias Mann from Max Planck Institute of Biochemistry for continuous support. We thank Tamara Díaz and Katharina Zettl (MPIB) for their excellent technical assistance and MSc. Mauro Rosales for an excellent article review.

## 6. Conflict of interest statement

The authors declare no conflict of interest in publishing these data. These data have not been published in any other scientific journal before.

## 7. Funding

This work was supported by the German Ministry of Education and Science (01DN18015) and the Max-Planck Society for the Advancement of Science. This work was also supported by the Center for Genetic Engineering and Biotechnology, La Habana, Cuba.

## 8. Author Contributions

All authors have read and agreed to the published version of the manuscript. Their contribution is stated. Conceptualization, D.V-B; V.B; L.J.G; I.B-R; Methodology, D.V-B; V.B; Y.R; J.R.W; Formal analysis, D.V-B; V.B; A.H-S; S.B; S.P; O.G; Writing—original draft preparation, D.V-B; V.B; A.H-S; S.B; Writing/review and editing, J.R.W; L.J.G; I.B-R; Project administration, J.R.W; L.J.G; I.B-R.

## 9. Data Availability

All data of proteomic and phosphoproteomic results generated during this study are included in this published article and supplementary information files. Any additional request would be available from the corresponding author on reasonable request.

## 12. Supplementary Information

**Fig 1S.**
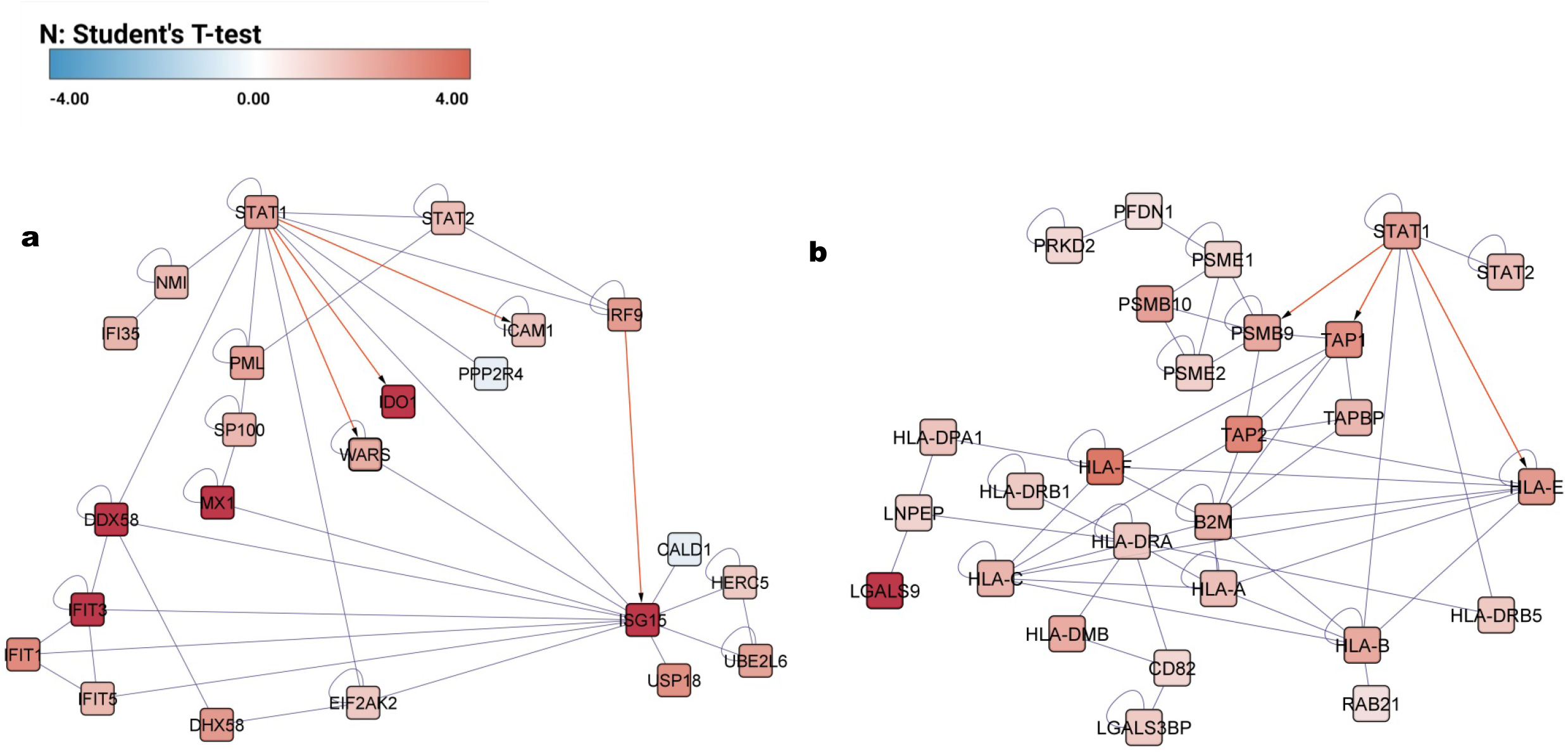
Network representation of the interconnected elements in the proteome data from STAT1/STAT2. a: components of the antiviral response, b: components of the MHC and proteasome. Color intensities represent protein changes in treated samples compared to cell control, as in the bar intensity color according to Student’s T-test. Bisogenet (v.3.0.0) plugin was used on the Cytoscape framework (v.3.8.2).

**Fig 2S.**
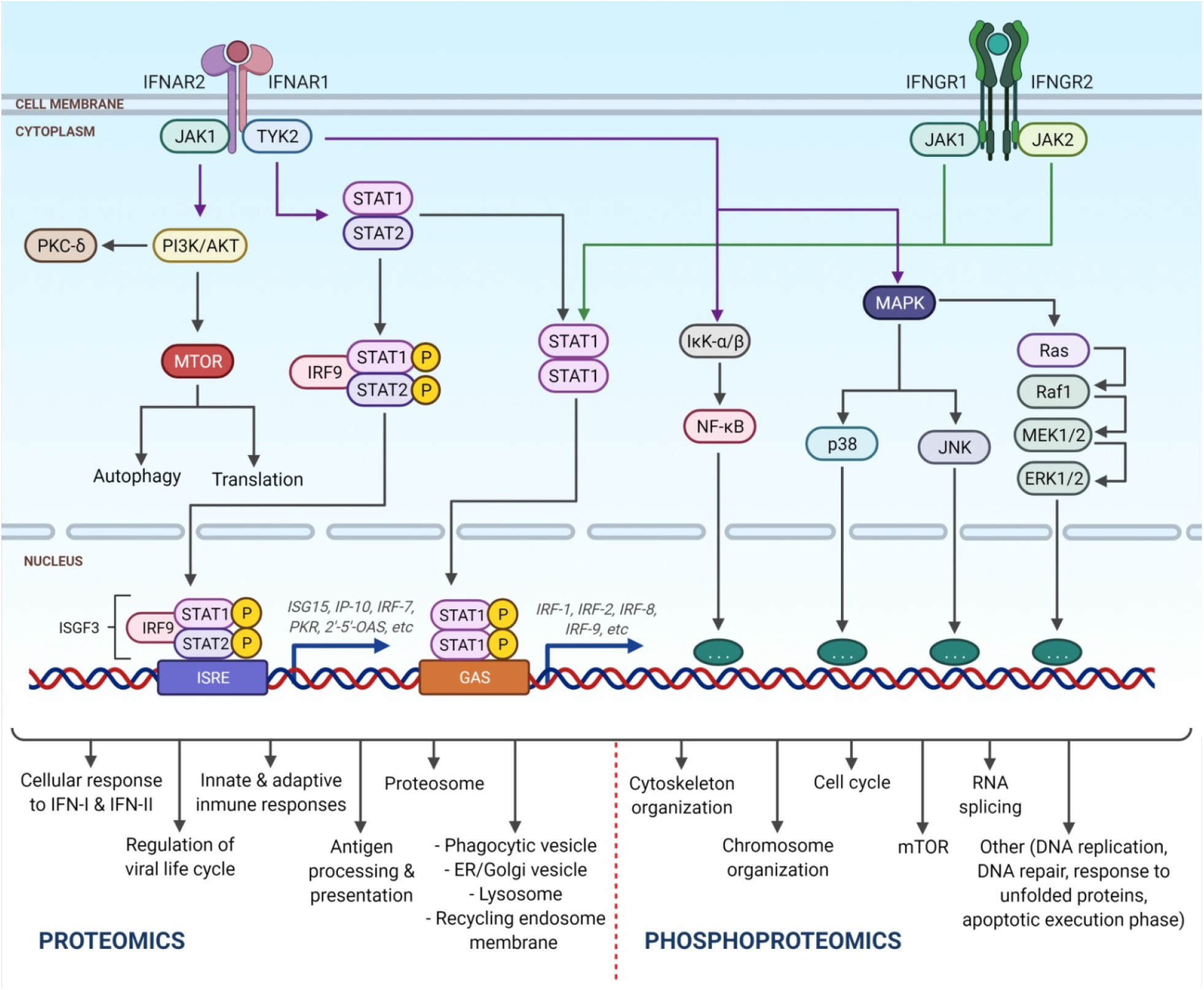
General pathway and biological processes regulated by HeberFERON in U-87 MG as it is shown in proteomic and phosphoproteomic data. The figure was created with BioRender (biorender.com).

**Table 1S. Results of Phosphoproteomic experiment**. Sequence window, Protein, and Gene names, Amino acid, and Position, P:-Log Student’s T-test p-value Intensity HF_Intensity CC, Student’s T-test Difference Intensity HF_Intensity CC and Student’s T-test Test statistic Intensity HF_Intensity CC.

**Table 2S. Results of a proteomic experiment**. Unique peptides, Sequence coverage [%], P: - Log Student’s T-test p-value LFQ intensity HF_LFQ intensity CC, Student’s T-test Difference LFQ intensity HF_LFQ intensity CC, Student’s T-test Test statistic LFQ intensity HF_LFQ intensity CC, Majority protein IDs, Protein names, Gene names.

**Table 3S. Enrichment analysis of Pathways in proteomic data using Toppfun tool**. Pathways Toppfun, p-value, Genes Symbols.

**Table 4S. Analysis of kinase motifs using Motif-All algorithm in Phosphosite Plus**. Motif, ZScore, P-Value, Fold Change, Foreground Matches, Foreground Size, Background Matches, Background Size, Kinases.

## Notes

### Competing Interest Statement

The authors have declared no competing interest.

